# Oral delivery of the amylin receptor agonist pramlintide

**DOI:** 10.1101/2023.11.09.566408

**Authors:** Celimar Sinézia, Tháyna Sisnande, Luis Peña Icart, Luís Maurício T. R. Lima

## Abstract

**Aim:** The aims of this study were to design and characterize polymeric microparticles for oral delivery of pramlintide, a triple proline human amylin analogue clinically proved efficient in diabetes therapy.

**Methods:** The microparticles were prepared with gastric-resistant polymer Eudragit S100 by double-emulsion and solvent evaporation technique. The study responses were repeatability, encapsulation efficiency, yield, morphology, particle size, response to acidic and alkaline milieu and pharmacokinetics.

**Results:** We obtained spherical microcapsules, with particle size of 66 μm ± 11, with 83.2 % ± 2.7 efficiency for pramlintide entrapment and 67.6 % ± 2.1 yield. Intra-venous pramlintide free in solution showed a plasmatic half-life of 6.8 min in mice. In contrast, oral delivery of acid-resistant pramlintide-loaded microparticles in mice showed a protracted release for 120 min compared to 30 min obtained for pramlintide in solution.

**Conclusions:** Oral route is an alternative for therapeutic development of pramlintide formulations.

**Graphical abstract:** 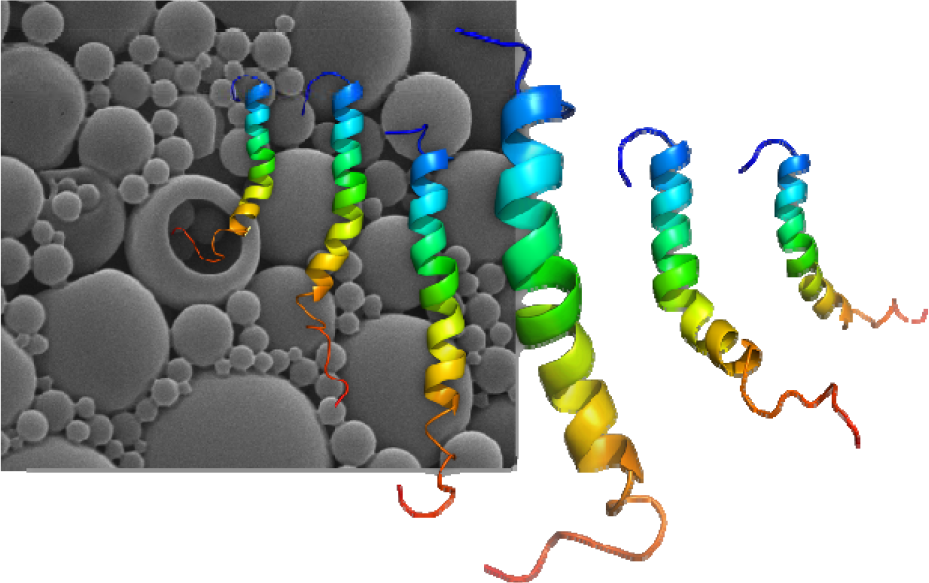

## 1. Introduction

The pancreatic β-cell may respond to varying pathophysiologic status, resulting in increased function during the build-up of metabolic resistance and subclinical diabetes, and converging to a decreasing function in overt diabetes outcome [1]. Diabetes is the result of failure in production, secretion and/or action of both insulin and amylin, both in post-prandial and fasting (basal) conditions. While insulin is essential for life, amylin is essential for a proper control of glucagon, satiety, gastric emptying, glycemic excursions, and other metabolic roles, also allowing expressive reduction in the insulin dose [2–4]. The restoration of complete pancreatic physiology in diabetic patients could only be achieved by simultaneous reposition of both β-cell hormones.

Since the discovery of amylin (also known as islet amyloid polypeptide, IAPP) [5, 6], its therapeutical use was immediately envisioned (patent US 5,367,052) [7] but it was not possible once amylin promptly aggregates into amyloid fibril in a broad pH range [8–10]. Proline-rich amylin in other species was found to be related to a more soluble peptide [11], inspiring the design of the triple proline human amylin variant ^[25,28,29]^Pro-hAmylin, named pramlintide (from **pr**oline **amlintide** = amilyn peptide) (AC-0137; CAS # 151126-32-8; patent US 5,998,367) [12]. However, pramlintide is also prone to amyloid aggregation in solution at a broad range of pH from acidic to alkaline [8, 13, 14], delaying the design of the product and entering the US market as first-in-class in 14 years after its first patent. Symlin® is the injectable pramlintide at pH 4.0 in sodium acetate solution approved by the FDA in 2005 [4]. The acidic Symlin® formulation is pharmaceutically incompatible with neutral to slightly alkaline insulins on the market, limiting its co-formulation with insulin [14–16] and leading to lack of interest in Symlin® besides its well-recognized benefits.

Several attempts have been developed in order to stabilize amylin and analogues, such as proteins, lipids, zinc and phytochemicals [14, 17–25], physico-chemical variables [19, 26–31], and conjugation with soluble polymers [29, 32, 33].

Oral delivery of peptides has long been pursued [34], and recently demonstrated clinically in humans for insulin [35–37], and the GLP-1 analogue semaglutide [38]. In this context, oral route pramlintide compatibility was considered a natural choice. Here we report the design, formulation and *in vivo* evaluation of gastric-resistant polymeric microparticles of Edragit® S100 loaded with pramlintide.

## 2. Material and Methods

### Material

Carboxy-amidated, C2-C7 dissulfide bond, pramlintide (TFA salt, lot #89437) was obtained from Genemed Biotech Inc (CA, USA). Chromatography-grade solvents were obtained from Tedia Brazil. All other reagents were of analytical grade.

### Ethics

The animal protocol was approved by the Institutional Bioethics Committee of Animal Care and Experimentation at the UFRJ (CEUA-CCS, UFRJ, #165/2019).

### Pramlintide-loaded Eudragit® S100 microparticles

Pramlintide (10 mg/mL in 5 mM sodium acetate pH 4.0; 200 μL) was added into the Eudragit® S100 solution (6 % w/v, in CH_2_Cl_2_:ethanol:isopropyl alcohol, 5:6:4, v/v/v; 5.0 mL) under agitation with Turrax™ T10 and dispersion probe S10N-10G (IKA) at 20,000 rpm for 10 min at room temperature. The resulting water:oil emulsion (W_1_/O) was added into 20 mL of aqueous polyvinyl alcohol (2% w/v; external aqueous phase) with agitation using Turrax™ T25 and dispersion element S25N-10G at 20,000 rpm for 10 min at room temperature. The second emulsion of water:oil:water (W_1_:O:W_2_) type was left under mechanical stirring for 16 h at room temperature in a fume hood for the removal of organic solvents. The microspheres were spun down by centrifugation (5,000 rpm for 15 min at 4 °C). The supernatant was removed and stored at -20 °C for further peptide quantification, using the fluorescamine method [39, 40], with six-points analytical curve in the range of 0.7 μg/mL to 50 μg/mL (linear regression R^2^=0.9991, properly corrected for blank). The microparticles were washed with water and centrifuged again, and this step was repeated two more times. The resulting material was flash-freeze with liquid nitrogen and lyophilized for three days without other additives. Dummy microparticles without peptide were formulated similarly, using as internal aqueous phase only 5 mM sodium acetate pH 4.0. A summary of the procedure is presented in **Table 1**

**Table 1.**
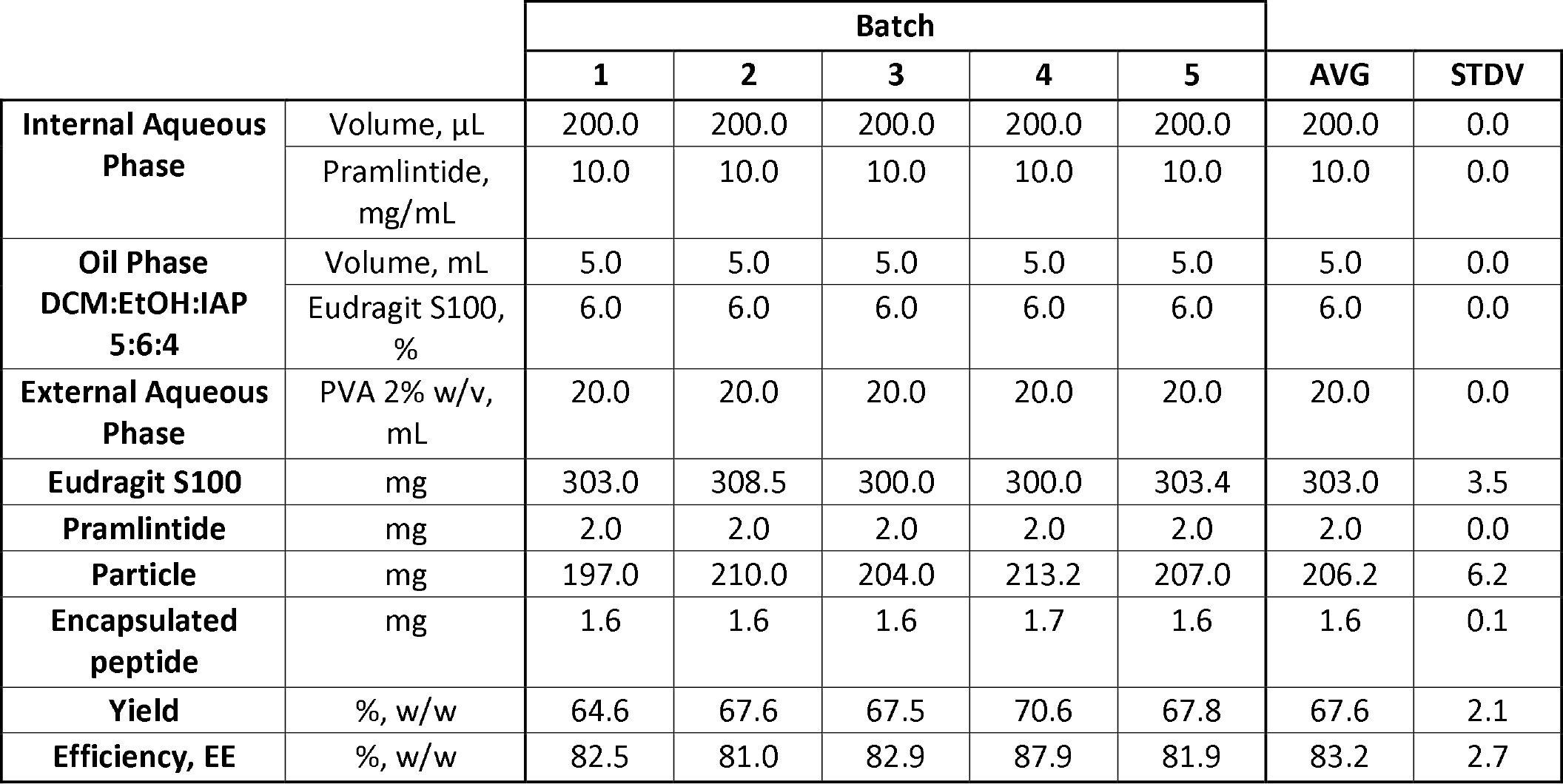
Preparation of Pramlintide-loaded Eudragit® S100 microparticles. Five batches of microparticles were prepared according to optimized method and the results are presented. Abbreviations: AVG, average; STDV, standard deviation of the mean; IAP, Internal Aqueous Phase; PVA, polyvinyl alcohol, DCM, CH_2_Cl_2_.

|  |  | Batch |  |  |  |  | AVG | STDV |
| --- | --- | --- | --- | --- | --- | --- | --- | --- |
|  |  | 1 | 2 | 3 | 4 | 5 |  |  |
| **Internal Aqueous Phase** | Volume, μL | 200.0 | 200.0 | 200.0 | 200.0 | 200.0 | 200.0 | 0.0 |
|  | Pramlintide, mg/mL | 10.0 | 10.0 | 10.0 | 10.0 | 10.0 | 10.0 | 0.0 |
| **Oil Phase**<br>**DCM:EtOH:IAP**<br>**5:6:4** | Volume, mL | 5.0 | 5.0 | 5.0 | 5.0 | 5.0 | 5.0 | 0.0 |
|  | Eudragit S100, % | 6.0 | 6.0 | 6.0 | 6.0 | 6.0 | 6.0 | 0.0 |
| **External Aqueous Phase** | PVA 2% w/v, mL | 20.0 | 20.0 | 20.0 | 20.0 | 20.0 | 20.0 | 0.0 |
| **Eudragit S100** | mg | 303.0 | 308.5 | 300.0 | 300.0 | 303.4 | 303.0 | 3.5 |
| **Pramlintide** | mg | 2.0 | 2.0 | 2.0 | 2.0 | 2.0 | 2.0 | 0.0 |
| **Particle** | mg | 197.0 | 210.0 | 204.0 | 213.2 | 207.0 | 206.2 | 6.2 |
| **Encapsulated peptide** | mg | 1.6 | 1.6 | 1.6 | 1.7 | 1.6 | 1.6 | 0.1 |
| **Yield** | %, w/w | 64.6 | 67.6 | 67.5 | 70.6 | 67.8 | 67.6 | 2.1 |
| **Efficiency, EE** | %, w/w | 82.5 | 81.0 | 82.9 | 87.9 | 81.9 | 83.2 | 2.7 |

### Efficiency and yield

The entrapment efficiency (EE) of pramlintide encapsulation was obtained by difference between the added (Pram_add_) and the remaining peptide in the supernatant (Pram_Sn_) from the first separation by centrifugation as follow:

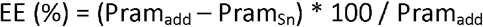

The encapsulation yield was obtained by the recovered mass of microparticles (MP_total_) in relation to the sum of initial polymer (Pol_add_) and peptide (Pram_add_) mass, as follow:

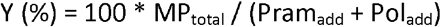

### Scanning electron microscopy (SEM)

The morphology of the microparticles was evaluated by SEM. The dry microparticles were dispersed over a carbon tape fixed onto a metallic stub. The material was coated with gold and the SEM were obtained in a FEI-Quanta 259 microscope operating at 12.5 kV with tungsten filament.

### Particle size distribution (PSD)

The particle size distribution of the Eudragit® S100 microparticles were evaluated by in a *Mastersizer 2000 – laser diffraction particle size analyser* (Malvern; Engepol-PEQ/COPPE/UFRJ). Microparticles (5 mg/mL) were dispersed in 50 mM sodium phosphate buffer pH 4.5 or pH 8.0 and kept under mild continuous magnetic stirring at room temperature and at given time intervals samples were subjected to PSD as indicated in the result section.

### In vivo pharmacokinetics

Adult Swiss male mice (20 g to 25 g) were randomized into three groups, with n=3 per time point:

G1 = intravenous pramlintide (5 μg pramlintide in 50 μL saline);

G2 = oral solution pramlintide (100 μg pramlintide in 200 μL saline);

G3–microparticle-entrapped pramlintide (100 μg peptide in 200 μL saline)

For oral administration, a curved stainless steel gavage cannula (22-gauge) was used. The animal was manually immobilized and the cannula inserted gently into the oral cavity and guided through the esophagus to the stomach. The volume of oral administration did not exceed the proportion of 10 μl per gram of body weight, respecting the limitation of the anatomical site. For intravenous administration, the animal was contained with the aid of a containment tube allowing free access to the animal’s tail. With the aid of a hypodermic needle (26-gauge, 13 mm x 0.45mm), the solution was administered into the lateral vein of the tail of the mouse, respecting the maximum administration volume of 200 μL per animal. At the end of the time for each subgroup, the animal was anesthetized and cardiac puncture was be performed. For anesthesia, intraperitoneal administration of ketamine/xylazine (80-100/10 mg/kg, intraperitoneal) was be performed. Eyelid and motor reflexes were evaluated to ensure complete state of anesthesia. Cardiac puncture was performed by inserting a needle (26-gauge) into the left atrium, collecting the largest possible blood volume. Euthanasia was performed with the administration of a lethal dose of anesthetic (100 mg ketamine / kg body weight; 2mg xylazine/ kg body weight) with intraperitoneal administration.

At given time intervals after administration (15min, 30 min, 60 min and 120 min for G1 and G2, and 15min, 60 min, 120 min, 180 min and 360 min for G3) the animals were anesthetized and the blood collected with EDTA by cardiac puncture, followed by centrifugation for separation of plasma which was stored at -20 °C for further analysis.

Pramlintide was quantified by ELISA (Merck Millipore, cat # EZHA-52) according to manufacturer instructions. The sensitivity for pramlintide was similar to human amylin as inferred from analytical curves from control.

## 3. Results

Pramlintide-loaded microparticles of Eudragit® S100 were prepared by the double emulsion and evaporation technique stabilized by PVA. The pramlintide encapsulation efficiency was 83.2 ± 2.7 % w/w and the microparticle yield was 67.6 ± 2.1 % w/w (n=5).

The resulting polymeric microparticles were spherical with smooth surface as observed by scanning electron microscopy (**Figure 1**).

**Figure.**
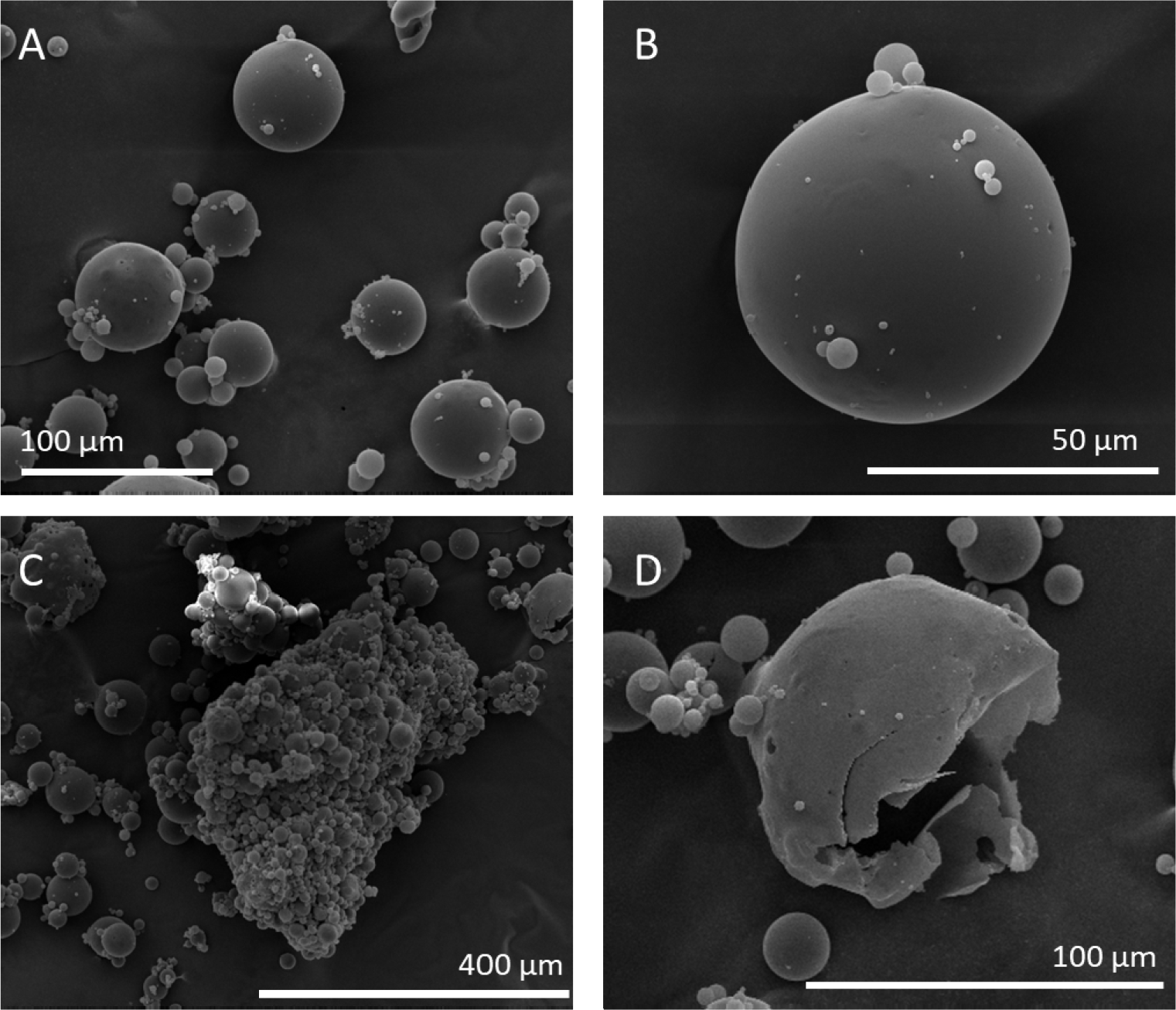
Particle morphology. The particle morphology was evaluated by scanning electron microscopy of the pramlintide-loaded Eudragit® S100 microparticles, evidencing A) spherical particles; B) uniform surface; C) small particles clumps; D) evidence for capsule (“hollow” sphere).

When dispersed in phosphate buffer pH 4.5, the particles showed a bimodal size distribution with populations at about 15 μm and 70 μm (**Fig. 2**), which persisted after two hours at this condition with a slight drift to higher sizes possibly by the formation of larger agglomerates (**Fig. 1C**) and a decreased repulsion between protonated carboxylic groups of the methacrylic acid of the polymer.

**Figure 2.**
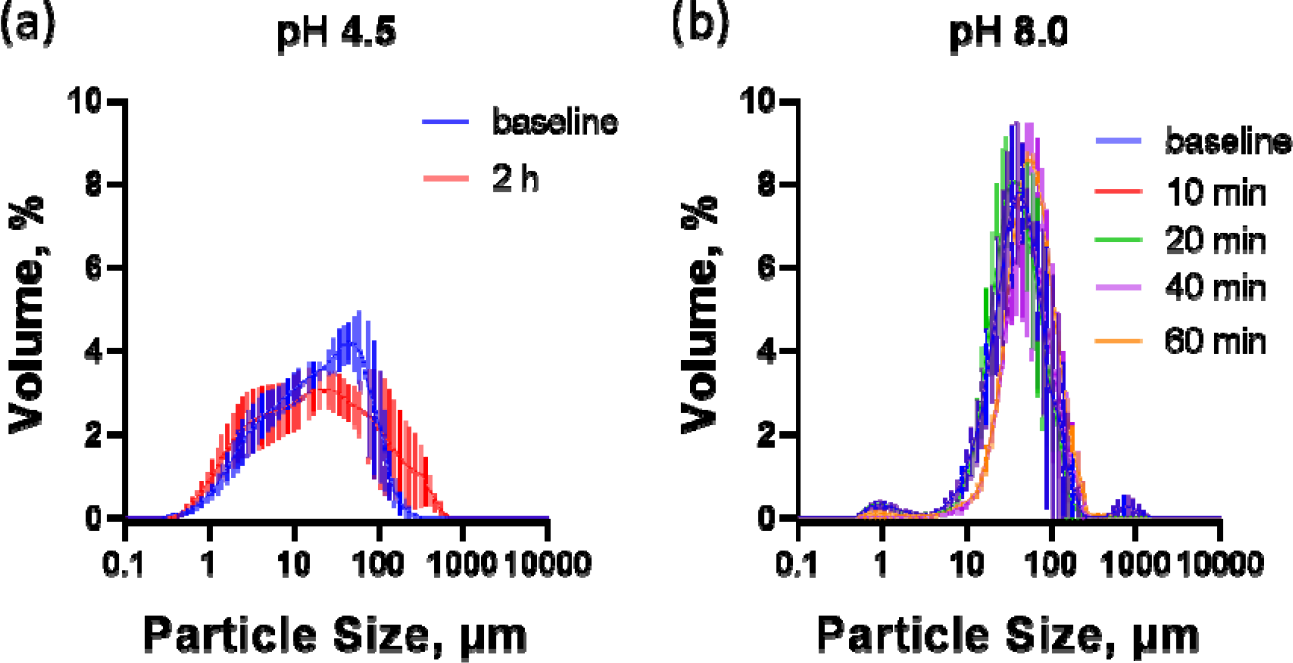
Particle size. The particle size distribution at acidic and mild-alkaline milieu was evaluated by laser scattering. The Eudragit® S100 microparticles were evaluated in (A) acidic solution (phosphate pH 4.5) up to 2 h, and (B) alkaline solution (phosphate pH 8.0) up to 60 min, at varying time intervals as indicated. Further attempts to measure were unsuccessful due to the lack of obscurity associated with dissolution of the polymer. Vertical bars represent standard error.

At pH 8.0 the particle distribution was unimodal at 66 ± 11 μm (**Fig. 2**). This behavior was maintained up to 60 min after dispersion in the phosphate buffer (66 ± 11 μm at 10 min, 70 ± 10 μm at 20 min, 75 ± 5 μm at 40 min, 74 ± 6 μm at 60 min). Further attempts to measure size distribution was not possible since the material was apparently solubilized, lacking sufficient obscurity for DLS measurements. These behavior at pH 4.5 and pH 8.0 confirms the desired pH dependence for enteric release.

### Pharmacokinetics *in vivo*

The release of pramlintide *in vivo* was evaluated comparing the peptide entrapped in the polymeric microparticles (Pram-MP) against the free peptide in solution administrated by intravenous and oral routes. Oral administration of Pram-MP was performed with a dose of 12.5 mg MP at 8 μg pramlintide/mg MP, corresponding to 100 μg pramlintide. The IV pramlintide (5 μg pramlintide /animal) resulted in a peak at about 3,700 pM at 15 min, and an exponential decrease with a half-life of 6.8 min.

The oral administration of free pramlintide in saline resulted in a fast absorption with a peak of 130 pM at 30 min, decreasing close to basal concentrations at 120 min. The pramlintide-loaded microparticles displayed a release with maximum at 180 min after about 120 min delay compared to oral solution pramlintide, indicating the expected behavior for the enteric polymer used.

## 4. Discussion

In this study we demonstrated the potential oral release of pramlintide, both free in solution and loaded into polymeric particles for enteric release. For the delayed enteric release, we formulated pramlintide into polymeric microparticles formulated with the gastric-resistant polymer Eudragit® S100 using the double emulsion technique

The resulting microparticles were uniform and spherical of about 50 μm diameter with smooth surface, which remain stable at acidic pH for up to 2h and at mild alkaline milieu for up to 40 min (**Fig. 1** and **Fig. 2**). The bimodal particle size distribution at acidic pH was most likely due to protonation of the carboxylic groups of the polymer resulting in decreased electrostatic repulsion and promoting agglomeration [41], which was not observed at pH 8.0 (**Fig. 2**).

The present microparticle formulation was initially based in preliminary studies with Eudragit® and the highly soluble peptide insulin [42, 43]. In our method we used PVA in the stabilization of the polymer-peptide emulsion, obtaining a high entrapment (83% w/w) and a reasonable yield (67% w/w), which we believe that can be possible to optimize by changes in buffers pH, solvent proportions in the emulsification, in wash step and mixing method.

Amylin or pramlintide administrated IV (10 μg) was reported to result in 10,000 pM plasmatic concentration in rats, while 1 μg pramlintide IV resulted in 1,000 pM plasma concentration [44]. The oral administration of soluble pramlintide resulted in a fast absorption with Tmax of 30 mi, reaching 130 pM which decreased to basal levels at 120 min, which is show to vary from 4 to 8 pM (**Fig. 3**). The absorption rate and extension after oral administration of peptide can vary according to anatomic and physiologic aspects, and the decrease in the absorption variability can be minimized by proper development of dosage forms, resulting in a predictable and trustable pharmacologic therapy [45].

**Figure 3.**
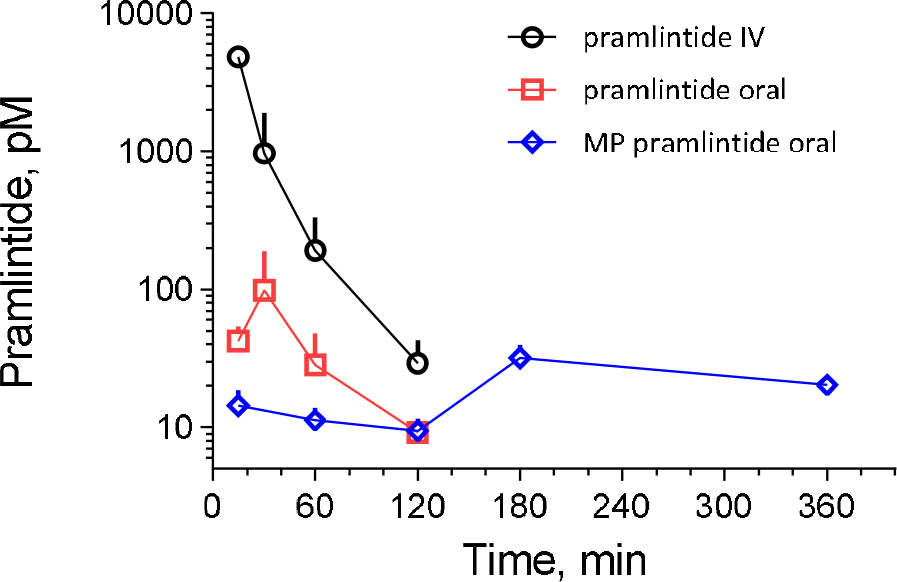
Pharmacokinetics of *in vivo* release of pramlintide. Pramlintide was administrated to mice (n=3 per time interval) by varying routes: intravenous (black circles; 0.2 μg/animal), oral free in solution (red squares; 4 μg/animal) and oral as microparticles (blue lozenges; 4 μg/animal). The available pramlintide in the circulation was evaluated by ELISA

We choose pramlintide since it is designed to be well stable at acidic acetic solution. However, we have recently demonstrated that human amylin can also be stabilized at acidic acetic milieu for hours [10], which could allow further development of the enteric microparticles for release of human amylin. Nonetheless, although homologous hormone reposition seems ideal, the use of the analogue pramlintide is favored since it has long been studied in clinical trials and approved for human use since 2005. In the present case we demonstrated that pramlintide can be delivered orally both for fast release from solution and for protracted release from microparticles. Further developments should optimize the formulation yields, extension of release as well as characterize both the proteoform products in the circulation and the pharmacologic activity, according to the International Conference of Harmonization (ICH) regulation. We have shown two dissimilar pharmacokinetic profiles for pramlintide, both free in solution and entrapped into microspheres for enteral release, and thus further developments will require addressing these two profiles and deciding which one – if any, or perhaps both – could follow to higher Technology Readiness Level.

## 5. Conclusions

Polymeric entrapment of the amylin receptor agonist pramlintide into biocompatible microparticles can be obtained with high efficiency and repeatability. The oral delivery of pramlintide can provide two pharmacokinetic profiles, both fast as soluble pramlintide, and with a protracted delivery when entrapped into the gastric-resistant polymer Eudragit® S100. These results demonstrate the feasibility of oral delivery of pramlintide, and potentially other amylin analogues.

## Acknowledgments

We would like to thanks the *Centro Nacional de Biologia Estrutural e Bioimagem* (CENABIO, UFRJ) for the access to the microscopy facility, and Prof. José Carlos Pinto for access to the particle size analyser at *EngePol* (UFRJ).

## Data Availability

The data generated during and/or analyzed during the current study are available from the corresponding author on reasonable request.

## Authorship statements (CRediT roles)

**Celimar Sinézia -** Data curation; Formal analysis; Investigation; Methodology; Writing - review & editing

**Tháyna Sisnande –** Investigation; Methodology; Writing - review & editing

**Luis Peña Icart** - Investigation; Methodology; Writing - review & editing

**Luís Maurício T. R. Lima -** Conceptualization; Data curation; Formal analysis; Funding acquisition; Investigation; Methodology; Project administration; Supervision; Validation; Visualization; Roles/Writing - original draft; Writing - review & editing

